# A high-throughput nematode sensory assay reveals an inhibitory effect of ivermectin on parasite gustation

**DOI:** 10.1101/2023.04.25.538347

**Authors:** Leonardo R. Nunn, Terry D. Juang, David J. Beebe, Nicolas J. Wheeler, Mostafa Zamanian

## Abstract

Sensory pathways first elucidated in *Caenorhabditis elegans* are conserved across free-living and parasitic nematodes, even though each species responds to a diverse array of compounds. Most nematode sensory assays are performed by tallying observations of worm behavior on two-dimensional planes using agarose plates. These assays have been successful in the study of volatile sensation but are poorly suited for investigation of water-soluble gustation or parasitic nematodes without a free-living stage. In contrast, gustatory assays tend to be tedious, often limited to the manipulation of a single individual at a time. We have designed a nematode sensory assay using a microfluidics device that allows for the study of gustation in a 96-well, three-dimensional environment. This device is suited for free-living worms and parasitic worms that spend their lives in an aqueous environment, and we have used it to show that ivermectin inhibits the gustatory ability of vector-borne parasitic nematodes.

## Introduction

Living organisms must sense their environment to find food, reproduce, and avoid predators or pathogens. Neural systems are necessary for the detection of cues from the environment and integration of a behavioral response. *C. elegans* has been a powerful model for understanding the neural and genetic basis for these behaviors in multicellular eukaryotes, and the assays developed using *C. elegans* have been used for comparative physiology across the nematode phylum.

Assays first established 50 years ago continue to be potent, simple tools for understanding complex behaviors in many nematode clades (Dusenbery, 1973; Ward, 1973). These are generally performed on a two-dimensional plane created by an agarose plate that deters nematodes from burrowing. Agarose plates are used for a variety of sensory assays and have been used to describe behavioral responses to hundreds of chemical and mechanical cues, but they are particularly well suited for the study of volatile sensation (olfaction). The ease and simplicity of this chemosensory assay has led to olfaction by *C. elegans* being one of the most thoroughly studied and understood behavioral processes in the entire animal kingdom (Ferkey et al., 2021). On the other hand, assays probing worm sensation of water-soluble molecules (gustation) are more difficult and have lower throughput, limiting the number of strains and cues that can be screened in parallel.

The relevance of olfaction to parasitic nematodes, which have stages that are entirely host-contained in aqueous environments, is unclear. Parasites such as vector-borne filarial nematodes, which are distantly related to *C. elegans* and cause devastating diseases like lymphatic filariasis and river blindness, among others, spend the vast majority of their life cycle in either an arthropod vector or a mammalian definitive host, with only an infinitesimal time spent on the host skin before being wicked into the puncture wound left by a feeding vector. For these nematodes, it is likely that gustation is more prominent than olfaction, yet the assays used to probe sensation of vector-borne nematodes are almost identical to those first developed using *C. elegans*. These assays have reliably shown that filarial nematodes respond to a variety of host cues (Fraser et al., 2018; Kusaba et al., 2008; Mitsui et al., 2021; Wheeler et al., 2020), but the throughput is low and the quantitative output lacks sensitivity, relying on large effect sizes in order to statistically resolve behavioral differences.

Recently, three-dimensional hydrogel-based nematode burrowing assays have been developed, and it has been shown that rearing nematodes in a three-dimensional environment reveals natural behaviors that are masked when using two-dimensional arenas (Baiocchi et al., 2020; Guisnet et al., 2021). Additionally, microfluidic devices have been designed to probe nematode sensory behaviors in an aqueous environment, increasing quantitative sensitivity (Chronis et al., 2007; Hu et al., 2015). We sought to combine these concepts to develop an assay that examines nematodes in a three-dimensional aqueous environment, has higher throughput than classical and contemporary microfluidic and burrowing assays, and allows for comparative physiological approaches by being applicable to a diversity of nematode species.

This assay could also be leveraged to study drug action on nematodes. Anthelmintics are known to affect migration of parasite nematode larvae, and anthelmintic resistance can involve mutations associated with sensory structures. An assay that probes the ability of nematodes to migrate toward a cue when challenged with a drug will test not only the drug’s impact on neuromuscular coordination but also its neurosensory effects. Thus, new high-throughput assays that examine nematode gustation may offer a useful alternative for identifying anthelmintic compounds and anthelmintic resistance.

With these goals in mind, we adapted the Stacks device, which creates open, reconfigurable tissue culture environments to investigate multicellular interactions in three dimensions (Yu et al., 2019). Stacks facilitate the communication and migration of individual cell types, and we reasoned that a similar behavior could be elicited in free-living and parasitic nematodes. Here, we describe the design and optimization of an open microfluidic environment for examining nematode sensation, allowing for high-throughput sensory assays that can examine the behavior of thousands of nematodes at a time.

## Results and discussion

### Open microfluidic devices can be used for high-throughput quantification of *C. elegans* gustatory behavior

Nematode gustation has classically been investigated using the “drop test” or two-dimensional agarose plates prepared with a gradient of water-soluble molecules (Bargmann and Horvitz, 1991; Hilliard et al., 2002). These experimental designs have been fruitful but have limitations in throughput and do not examine the animal in a more natural three-dimensional environment. When hydrogels have been used to examine nematode behaviors in three dimensions, they have maintained the traditional approaches of using large-format dishes or wells (Baiocchi et al., 2020; Guisnet et al., 2021; Lesanpezeshki et al., 2019). We combined advances in microdevice fabrication and hydrogel utilization for nematode culture to design a 96-well device that allows nematodes to freely burrow through a three-dimensional environment toward a cue or away from a repellent.

Previously, Stacks devices had been used to examine cellular chemotaxis in a complex tissue culture environment (Yu et al., 2019). These devices can be reconfigured within a given experiment, allowing the removal or replacement of cells, nutrients, and/or cues that were present in individual slices of the stack. Initial attempts leveraging unmodified Stacks devices, which included 24 wells, were highly successful (Fig. 1). Adult wild type *C. elegans* showed attraction to 100 and 200 mM NaCl, with the chemotaxis index (CI) being slightly but insignificantly greater when the attractant was placed on top of the column than on the bottom (Fig. 1B). All remaining experiments set up the assay with cues on the top of the well.

**Figure 1.**
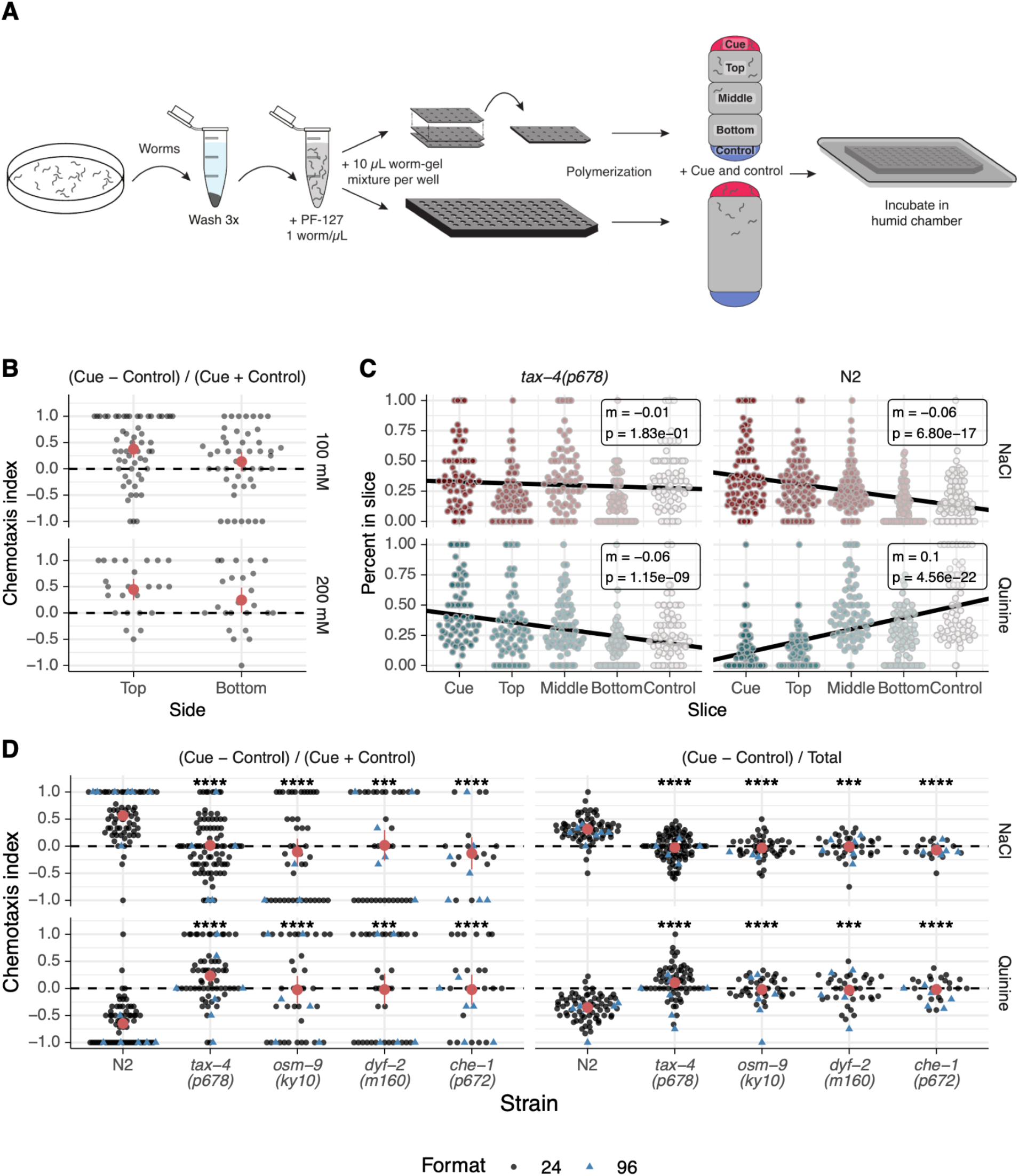
An open microfluidic device for high-throughput *C. elegans* gustatory assays. A schematic overview of the procedures used to produce the data in B-D. *C. elegans* are washed from NGM culture plates into a 1.5 mL tube and washed 3 times to remove residual debris and bacteria and to enrich for young adult worms. Worms are then mixed with liquid PF-127 and quickly added to the devices to allow for polymerization of the hydrogel. Initial experiments used 24-well devices with 3 slices, and later experiments used the gustatory microplate. Cues and controls are added as liquid bubbles to the top and bottom of the polymer, and then devices are incubated in a humid chamber to allow for worm gustation and burrowing. (B) *C. elegans* are attracted to either 100 mM or 200 mM NaCl, with a slightly higher CI when the cue was added to the top of the well. (C) Three-layer Stacks devices were separated after 1 hr. incubation, and worms in each slice were counted. Wild type (N2) worms are attracted to 200 mM NaCl and repelled by 10 mM quinine. *tax-4(p678)* worms show no response to NaCl and are attracted to quinine. (D) Chemotaxis indices of five chemosensory-defective knockout strains of *C. elegans* in response to 200 mM NaCl and 10 mM quinine. In each figure, dots represent the value for a single well of the device, large red dots represent the mean, and error bars represent the confidence limits. ***: p <= 0.001, ****: p <= 0.0001.

The devices allowed for worms to move either up and toward an attractant (i.e., NaCl) or down and away from a repellant (i.e., quinine), and knock-out strains known to be insensitive to salt attraction showed no response to NaCl (Fig. 1C). Using the reconfigurable 3-layer devices allowed us to separate the individual slices after incubation and count the number of worms in each slice. The number of wild type (N2) worms in each slice showed a remarkable linear relationship with the distance from the cue added at the top of the well, which resulted in a significant non-zero slope when the slices were numbered 1-5 (with slice 1 being the cue bubble and slice 5 being the control bubble, Fig. 1A). *tax-4(p678)* worms, which are insensitive to NaCl, had a slope that was not significantly different from zero (Fig. 1B). Interestingly, in these assays *tax-4(p678)* seemingly showed attraction to quinine -a response that to our knowledge has not been noted. These data demonstrate the unique quantitative sensitivity enabled by the Stacks devices.

The ability to reconfigure and count individual slices in the stack was useful, but for our experimental purposes we preferred increased throughput in contrast to increased customization. The fabrication of significantly larger reconfigurable Stacks devices was unlikely to achieve the required flatness (even some of the 24-well slices had to be discarded and carefully selected due to a slight bend in the slice), so we designed a second device that was not reconfigurable but included 96-wells and had the same dimensions as a 96-well microtiter plate (Fig. S1). We call this device the gustatory microplate. This new design allowed us to dramatically scale up the number of worms and strains that could be examined in parallel, allowing for multiple replicates of 5 strains and 2 cues to be completed in a matter of hours, showing that a range of knockout strains with sensory defects were unable to respond to NaCl or quinine (Fig. 1D). The data from 96-well devices did not differ from those acquired with 24-well reconfigurable devices. Increased replication allowed us to clearly show the significant positive CI of *tax-4(p678)* to 10 mM quinine (CI = 0.231, p = 0.0004; CI_Total_ = 0.106, p = 0.006). Previous experiments using the drop assay on NGM plates showed that *tax-4(p678)* retained a strong avoidance to quinine (Hilliard et al., 2004). The interesting behavioral response of *tax-4(p678)* to quinine in a three-dimensional environment is one that the gustatory microplate is uniquely poised to explore and confirm findings that three-dimensional environments reveal previously unknown natural behaviors (Guisnet et al., 2021).

Sensory defective strains of *C. elegans* in the gustatory microplate behaved similarly to those on traditional chemotaxis agarose plates. That is, worms tended not to roam through the matrix, but instead showed little directed movement at all. When we included a computational filter that required that more than two worms reach either the cue or control bubble, the sample size was substantially reduced while the conclusions and statistical significance of the differences between strains were not altered. *che-1(p672), osm-9(ky10)*, and *dyf-2(m160)* had the sample size reduced by 46%, 54%, and 79%, respectively (Fig. S2). Thus, we did not include such a filter in any data acquired with the gustatory microplate.

### The gustatory microplate recapitulates behavioral phenotypes of vector-borne parasitic nematodes while increasing throughput

A significant goal of the design of the gustatory microplate was broad applicability for nematodes with diverse life history traits. Most nematode behavioral assays were designed first with *C. elegans* in mind and then only translated for use with parasites that had similar life-history traits (i.e., free-living stages that are amenable to culture on agarose plates). We decided to take the opposite approach - start with a design that had a greater likelihood of being applicable for parasites, especially those that do not have free-living stages.

We and others have previously shown that the L3 stage of vector-borne filarial nematodes specifically respond to a variety of cues associated with mammalian skin and internal tissues (Fraser et al., 2018; Kusaba et al., 2008; Mitsui et al., 2012; Mitsui et al., 2018; Mitsui et al., 2021; Wheeler et al., 2020). These were performed in two-dimensions on agarose plates, upon which L3s show limited translational movement (parasite larvae are placed only a few millimeters away from the cue, in comparison to *C. elegans* being placed several centimeters away from a cue). In the 96-well gustatory microplate, the vector-borne parasitic nematode *Brugia pahangi* showed a significant positive attraction to serum (Fig. 2B). The CI in the device (CI = 0.75) was similar to that previously calculated using agarose plates (CI = 0.87 from (Wheeler et al., 2020) and CI = 0.9 from (Kusaba et al., 2008)). Importantly, the miniaturization of the assay allowed a single researcher to easily setup 2 microplates per experiment, totalling 192 wells, a more than ten-fold increase in throughput of one of the few behavioral assays available for parasitic stages of filarial nematodes. We anticipate that an experienced user will be able to set up and score >5 plates in a single experiment, with a bottleneck primarily arising from the time it takes to count worms in each well. We are currently optimizing high-content imaging of the gustatory microplate, which will dramatically increase the throughput.

**Figure 2.**
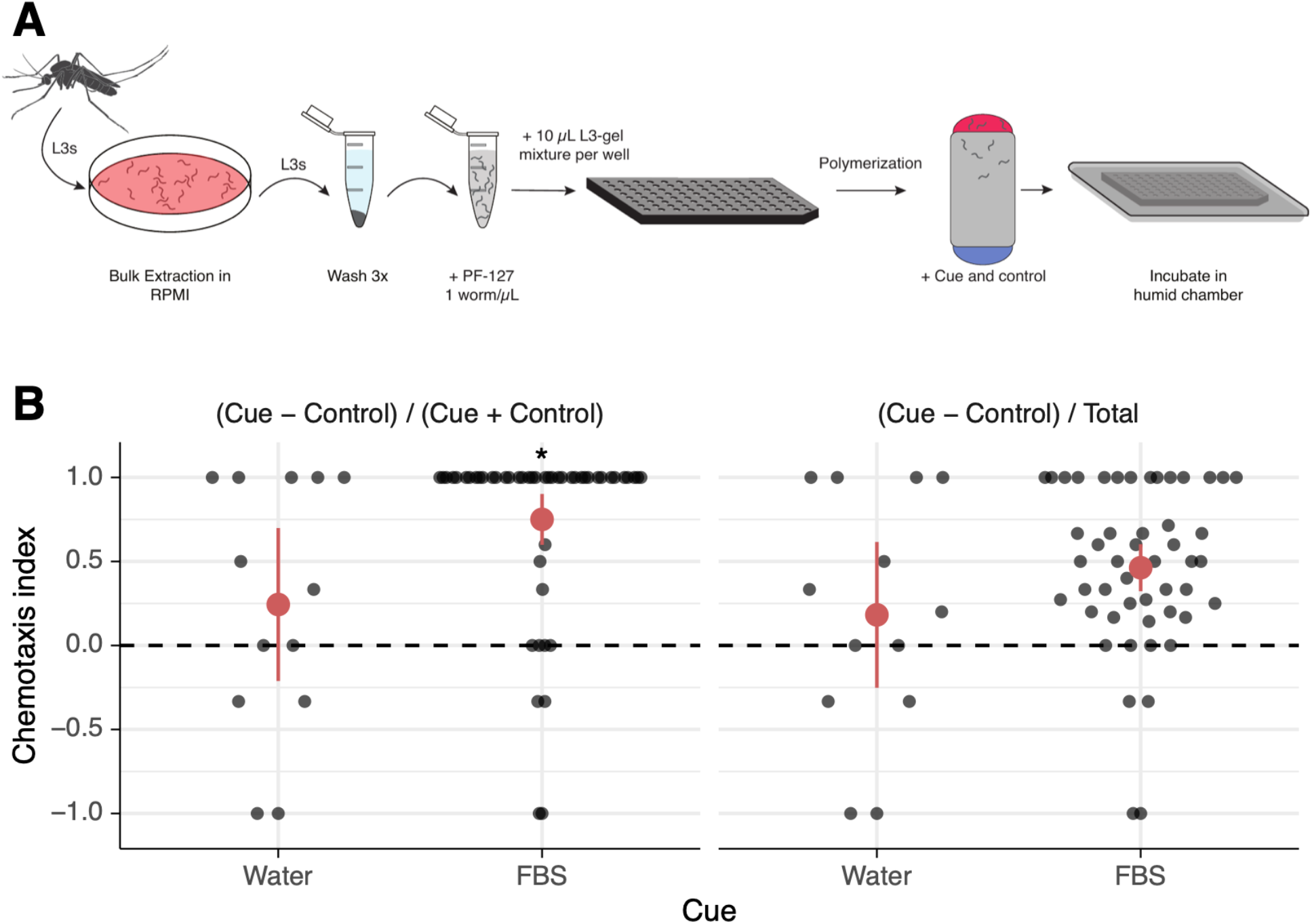
An open microfluidic device for high-throughput filarial nematode gustatory assays. (A) A schematic overview of the procedures used to produce the data in B. L3s were extracted from mosquitoes and washed 3 times prior to being mixed with liquid PF-127 and added to devices to allow for polymerization. Cues and controls are added as liquid bubbles to the top and bottom of the polymer, and devices are incubated in a humid chamber to allow for worm gustation and burrowing. (B) L3s from *Brugia pahangi* are attracted to FBS. Two methods for calculating chemotaxis index are shown. Dots represent the CI for a single well of the device, large red dots represent the mean, and error bars represent the confidence limits. *: p <= 0.05.

### Ivermectin reduces the gustatory ability of vector-borne parasitic nematodes

Ivermectin is a constituent of most anthelmintic treatments in mass drug administration against filariases like river blindness and lymphatic filariasis. In nematodes, ivermectin acts as an agonist of glutamate-gated chloride channels, but the mechanism of action and its effect on parasite behavior is incompletely understood. *C. elegans* strains that are resistant to ivermectin often have defects in their sensory capabilities due to abnormal development in sensory neurons (Dent et al., 2000; Page, 2018), and we reasoned that ivermectin might have a similar effect on parasite sensation.

The increased throughput allowed by the gustatory microplate enabled the analysis of the effects of ivermectin and albendazole, an anthelmintic that targets β-tubulin, on the sensory capabilities of filarial nematode larvae. Treatment with ivermectin, but not albendazole or DMSO, resulted in the inhibition of larval migration to FBS, and this was true for both *B. pahangi* and *Dirofilaria immitis*, the canine heartworm (Fig 3A-B). The CI for control worms was 0.84 and 0.59 for *B. pahangi* and *D. immitis*, respectively, while 100 µM of ivermectin reduced these values to 0.130 and 0.062 (Fig. 3B). While this concentration is much higher than that found in serum after ivermectin treatment *in vivo*, micromolar concentrations of ivermectin are regularly used to explore its effects on parasites *in vitro*, and the strong effects on gustation were seen after a short 20 minute treatment - far shorter than other *in vitro* assays that usually only see effects after >24 hrs (Harischandra et al., 2018; Loghry et al., 2020; Moreno et al., 2010). Importantly, inhibition of migration was not due to a generalized decrease in health or neuromuscular capabilities, as the proportion of *B. pahangi* L3 migrating to either the top or bottom of the device was not significantly changed by ivermectin treatment (Fig. 3C). Treatment of *D. immitis* with ivermectin did reduce the ability of L3s to migrate, but only by 10%, which is unlikely to completely explain the much larger decrease in CI.

**Figure 3.**
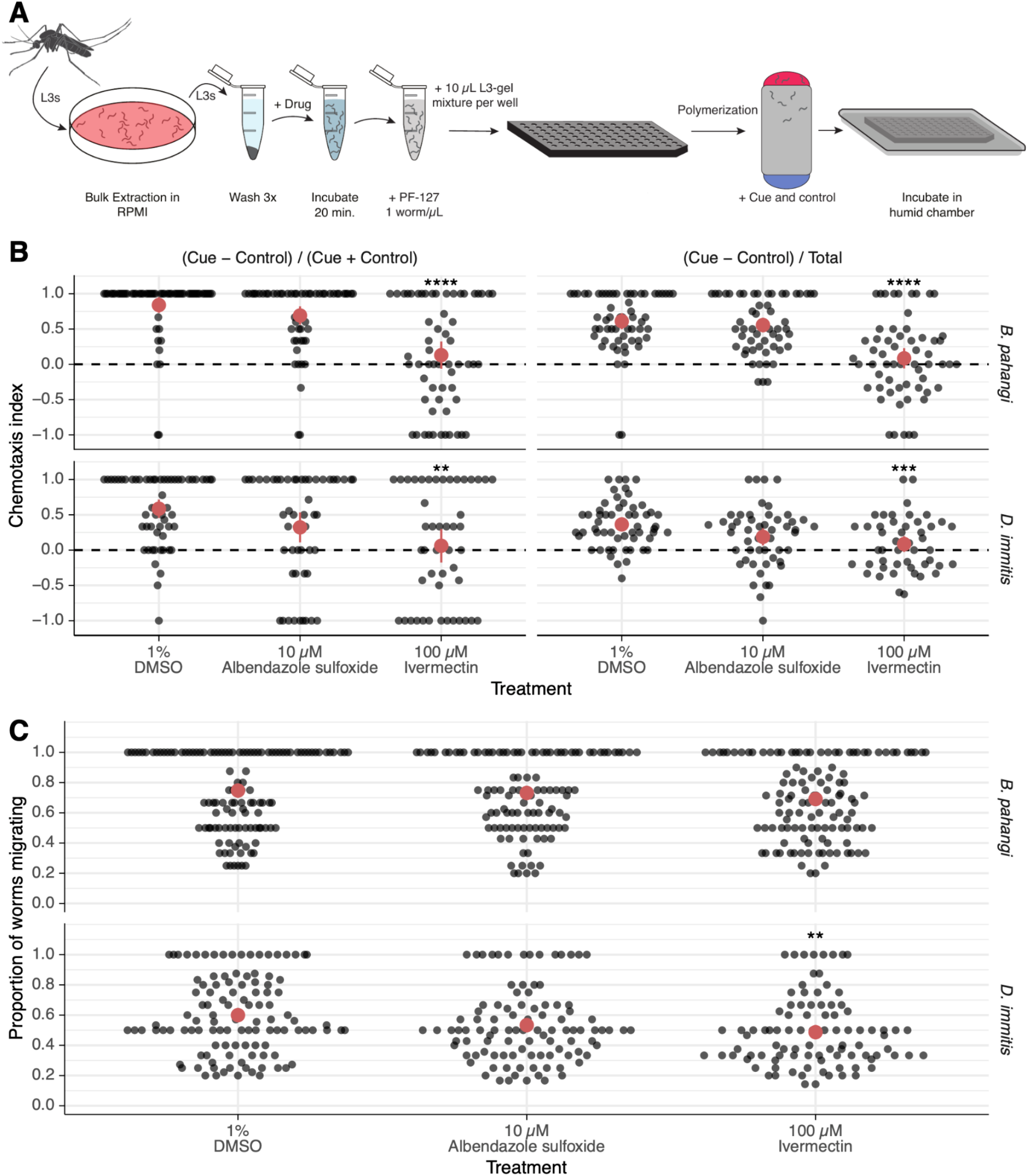
Ivermectin significantly inhibits vector-borne parasitic nematode gustation. (A) A schematic overview of the procedures used to produce the data in B-C. Worms and devices are prepared as in Fig. 2A, but a 20 minute drug treatment is performed prior to adding worms to devices. (B) 100 µM ivermectin, but not 1% DMSO or 10 µM albendazole sulfoxide, significantly inhibits migration toward FBS by L3s from either *Brugia pahangi* or *Dirofilaria immitis*. Two methods for calculating chemotaxis index are provided. (C) Similar proportions of control or treated worms move to either cue or control bubbles. 100 µM ivermectin slightly but significantly reduces the movement of *Dirofilaria immitis* L3s. In each figure, dots represent the value for a single well of the device, large red dots represent the mean, and error bars represent the confidence limits. **: p <= 0.01, ***: p <= 0.001, ****: p <= 0.0001.

Disrupting worm sensation is a potential route for parasite treatment and control, and mutations associated with nematode sensory structures are associated with anthelmintic resistance in *C. elegans* (Dent et al., 2000; Page, 2018; Wheeler et al., 2022). The mechanism of action of ivermectin against helminths is complex and likely varies based on the species and stage of the parasite it is acting on. To our knowledge, this is the first report of an anthelmintic having an effect on the sensory ability of parasitic nematodes.

High-throughput assays that recapitulate a three-dimensional environment will be important not just for comparative studies of nematode behavior, but also for the evaluation of anthelmintic resistance and identification of novel anthelmintics that combat this resistance. One assay using a three-dimensional arrangement - the larval migration assay (LMA) - has been developed to assess the drug sensitivity of some nematodes (Evans et al., 2017; Kotze et al., 2006). In this setup, treated larvae migrate down through agar and a filter into a receiver plate below; worms sensitive to the drug are unable to complete this process. The LMA has been used to identify compounds with anthelmintic properties and drug-resistant wild isolates (Nixon et al., 2019). Approaches such as the LMA are useful alternatives to standard screening regimens that measure gross motility in microtiter plates, and these will be made more powerful if they can reliably evaluate neurosensory and neuromuscular activity and coordination for a diverse set of nematodes.

Our modified gustatory microplates are positioned to enable new investigations into comparative nematode physiology and drug responses. Important advances have been made in understanding the divergent sensory profiles among Clade IV and V nematodes - clades including parasites such as threadworms and hookworms, which have free-living stages, and entirely free-living nematodes like *C. elegans* (Bryant and Hallem, 2018; Bryant et al., 2022; Castelletto et al., 2014; Gang et al., 2020). However, the free-living stages of these species readily crawl on the surface of an agarose plate, while many of the most important parasite species do not. Given the success of three different nematode species in gustatory microplates - free-living *C. elegans* and two species of vector-borne filarial nematodes - we anticipate broad applicability across the phylum.

## Materials and methods

### *C. elegans* strains and maintenance

N2, PR678, PR672, SP1234, and CX10 strains were obtained from the *Caenorhabditis* Genetics Center (CGC) and maintained at 20°C on NGM plates seeded with *Escherichia coli* OP50. Propagation of worms was maintained by routine picking of L4 stage worms to seeded NGM plates.

### Parasite maintenance

*Brugia pahangi* and *Dirofilaria immitis* microfilariae (mf) in blood were supplied by the NIH/NIAID Filariasis Research Reagent Resource Center (FR3 (Michalski et al., 2011)). Mf were fed in blood to female adult *Aedes aegypti* mosquitoes (LVP and SD strains for *B. pahangi* and *D. immitis*, respectively, at 90-120 mf/20 µL) and developed to the L3 stage for extraction. L3s were extracted into warm RPMI 1640 (Sigma) from mosquitoes on day 14 post-infection, as previously described (McCrea et al., 2021).

### Gustatory microplate design and fabrication

The 96-well gustatory microplate was modified from the original 24-well Stacks(Yu et al., 2019) to increase throughput. The device was designed to fit within a Nunc Omnitray single-well plate (ThermoFisher Scientific) and has a culture area of 3.14 mm^2^ and a working volume of 12.56 µL for each layer (Fig. S1). The device was then fabricated through standardized rapid prototyping methods using Micro-CNC milling techniques to fabricate each layer from sheets of polystyrene (4 mm, Goodfellow) using Autodesk’s Fusion 360 CAD software and a Tormach PCNC 770 Mill (Guckenberger et al., 2015). Following milling, the devices were cleaned using ultrasound sonication in isopropanol for 10 minutes to remove any debris or particles.

Custom holders were designed and fabricated using Fused Deposition Modeling (FDM) 3D printing on a 5th generation Makerbot, with white polylactic acid filament (Matterhackers) extruded using the Makerbot software’s coarse setting. The holders function as an aligner for securing Stacks layers on top of each other and as spacers to prevent device contamination when placed on a benchtop. The CAD files and dimensions for the gustatory microplate and custom holder are included (Supplementary Materials).

### Gustatory assays

In order to effectively study *C. elegans* in their natural three dimensional environment and increase the throughput of chemosensory assays, the gustatory microplates were used in combination with pluronic gel (Lesanpezeshki et al., 2019; Yu et al., 2019). To prepare *C. elegans* for this assay, five L4 worms were picked each onto 6 cm NGM plates 4-6 days prior to the assay date. In addition to this, a 30% w/w PF-127 (Sigma Aldrich)/water gel solution was created 2-3 days before assay date to allow the PF-127 to dissolve completely at 4°C.

On the day of the experiment, worms were added to liquid PF-127 to create a uniform worm-gel mixture. Mixed populations of worms were washed off of NGM plates with M9 and washed twice with M9 and once with water to remove bacteria and enrich for adult worms. Worms were counted, allowed to settle, and all but 50 µL of water was removed. 30% w/w PF-127 was added with a pre-cooled wide bore pipette tip and mixed gently to create a worm-gel mixture at 1 worm/µL with a final concentration of 26-28% PF-127. Worm-gel mixtures were briefly kept in ice water baths (12°C ± 1°C) until ready to add to the devices.

A single 96-well gustatory microplate or three 24-well Stacks bound tightly together were prepared for addition of worm-gel mixture by setting them in a device holder on a clean bench. A digital repeating pipette was pre-cooled by flushing with ice-cold water and then used to pipette 12 µL of worm-gel mixture into each well of the device. PF-127 will polymerize at room temperature in 5-10 minutes, during which worms were counted in each well. After polymerization, 3 µL of cue (200 mM NaCl for attraction assays and 10 mM quinine for aversion assays) was added to one side of each well and 3 µL control was added to the opposite side; these bubbles will diffuse through the polymer and keep the polymer column hydrated during incubation. Devices were placed in humid chambers and incubated for 1 hr. at room temperature, after which worms that migrated to the top and bottom side of the device were counted.

For assays with *B. pahangi* and *D. immitis* L3s, worms were isolated from infected mosquitoes 14 days post-infection via bulk extraction in RPMI 1640 at 37°C (McCrea et al., 2021). Thrashing L3s that emerged from mosquitoes during extraction were counted and manually transferred to a new microcentrifuge tube, in which they were washed three times with PBS and titered to a concentration of 1 L3/µL. For drug treatment experiments, the L3s were pre-treated for 20 min. at 37°C in 1% DMSO or various concentrations of albendazole or ivermectin. After treatment, 30% PF-127 was added and mixed gently to create a L3-gel mixture of 1 L3/µL. The same procedure for device preparation as described for *C. elegans* was used, and devices in humid chambers were incubated at 37°C for 1 hr. to allow for burrowing.

After incubation, worms in the liquid bubbles on each side of the well were counted using an upright Zeiss Stemi 305 dissecting microscope.

### Data analysis

Two forms of chemotaxis index (CI) were used and are annotated in each figure:

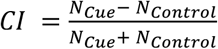, where N is the count in the relevant bubble after incubation.

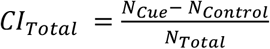, where N_Total_ is the count of all worms taken prior to incubation.

Only wells in which >0 worms migrated to the top or bottom liquid bubbles were used for all CI calculations. Requiring wells to include >2 worms moving to the top or bottom liquid bubbles greatly reduced the sample size without changing the statistical relationships (Fig. S2).

## Acknowledgements

Some *C. elegans* strains were provided by the CGC, which is funded by NIH Office of Research Infrastructure Programs (P40 OD010440). Parasite materials were provided by the NIH/NIAID Filariasis Research Reagent Resource Center (www.filariasiscenter.org).

## Author contributions

Conceptualization: NJW, MZ

Methodology: NJW, MZ, LRN, TDJ

Formal analysis: LRN, NJW

Investigation: LRN, NJW

Resources: MZ, DJB

Writing - original draft: LRN, TDJ, NJW

Writing - review & editing: LRN, TDJ, NJW, MZ, DJB

Visualization: NJW

Supervision: NJW, MZ, DJB

Project administration: NJW, MZ, DJB

Funding acquisition: MZ, DJB

## Data availability

All data and scripts for analysis and visualization are available at https://github.com/zamanianlab/ChemoDevice-ms.

---

**Supplementary Figure 1.**
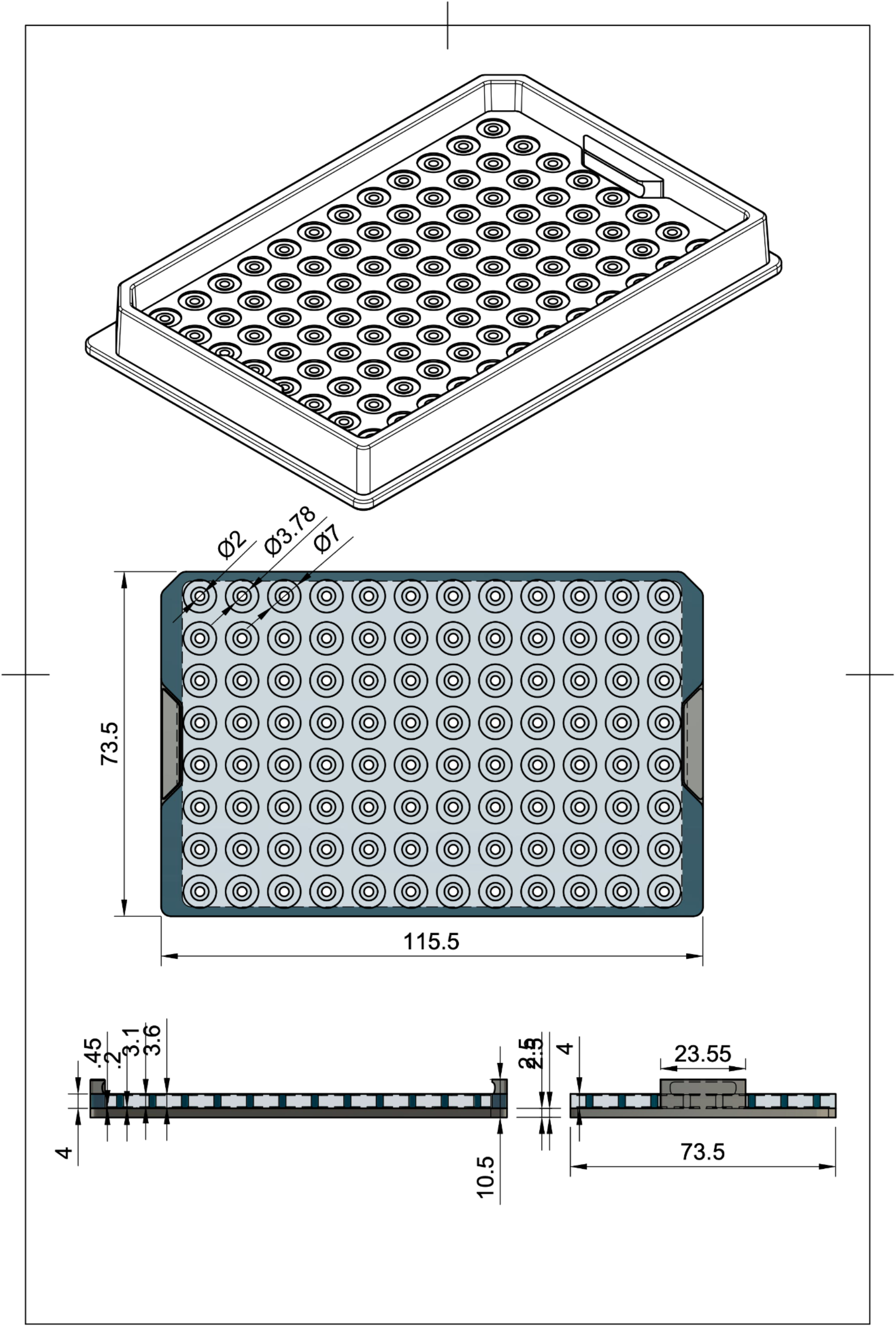
Model of the gustatory microplate and holder for experiment setup. Plans for milling and 3D printing the guy holder can be found at https://github.com/zamanianlab/ChemoDevice-ms.

**Supplementary Figure 2.**
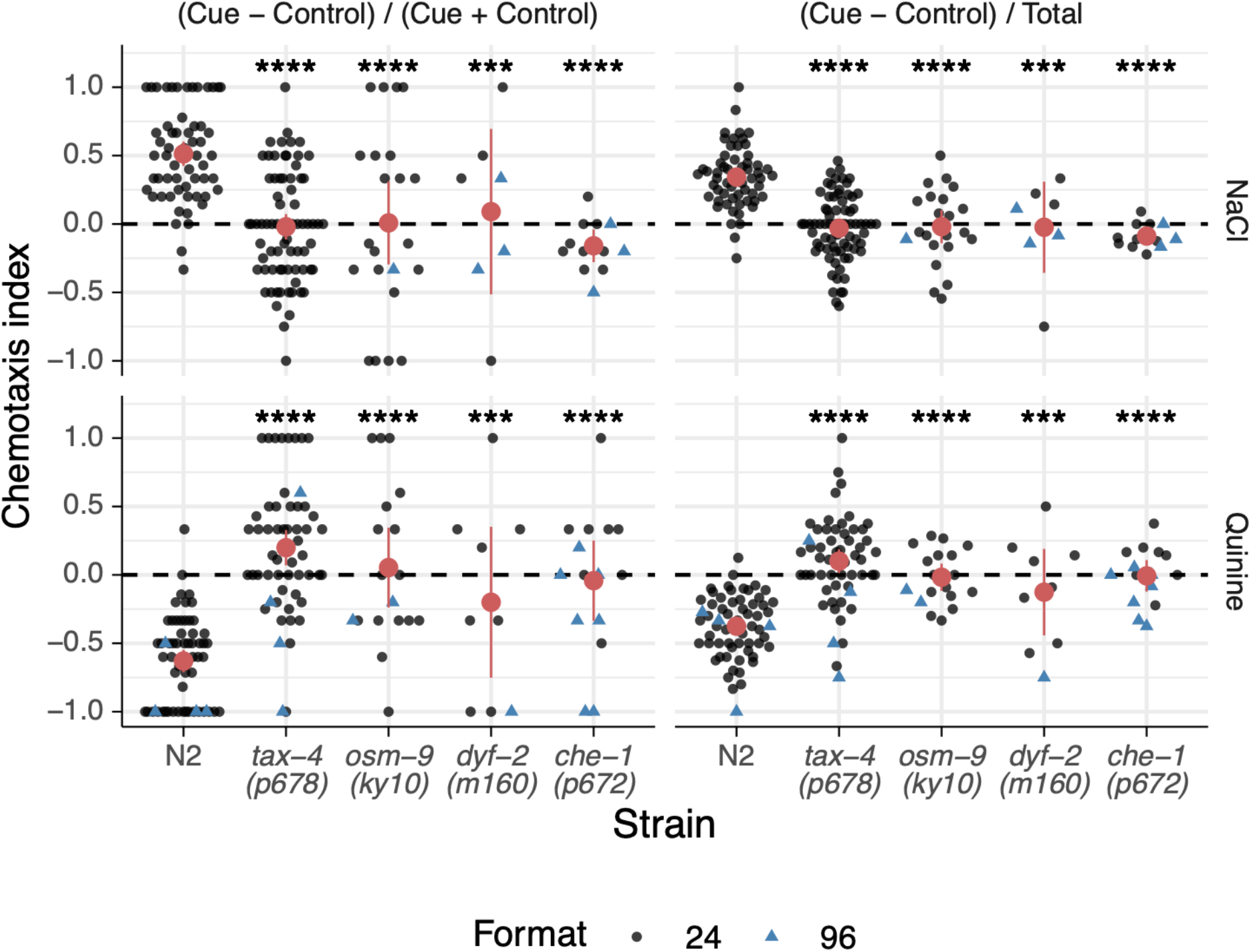
Data from Fig. 1D with a more stringent filter. Using only wells that had more than 2 worms reaching the cue or control bubble substantially reduced the sample size of the experiment.

